# Regressions Fit for Purpose: Models of Locust Phase State Must Not Conflate Morphology With Behaviour

**DOI:** 10.1101/174763

**Authors:** Swidbert R. Ott

## Abstract

Locusts are defined by their capacity to transform between two very distinct integrated phenotypes or ‘phases’ in response to changes in population density: a solitarious phase, which occurs when densities are low, and a gregarious phase, which arises as a consequence of crowding and can form very large and economically damaging swarms. The two phases differ fundamentally in their behaviour, physiology and morphology. A large body of work on the mechanistic basis of behavioural phase transitions has relied on multivariate logistic regression (LR) models to estimate the probability of behavioural gregariousness from multiple behavioural variables. Martín-Blázquez and Bakkali (2017, *Entomologia Experimentalis et Applicata* 163, 9–25) have recently proposed standardised LR models for estimating an overall ‘gregariousness level’ from a combination of behavioural and, unusually, morphometric variables. Here I develop a detailed argument to demonstrate that the premise of such an overall ‘gregariousness level’ is fundamentally flawed. Since locust phase transformations intrinsically entail a decoupling of behaviour and morphology, phase state cannot meaningfully be conflated onto a single axis. LR models that do so are therefore of very limited value for any analysis of phase transitions. I furthermore show why behavioural predictor variables should not be adjusted by measures of body size that themselves differ between phases. I discuss the models fitted by Martín-Blázquez and Bakkali (2017) to highlight potential pitfalls in statistical methodology that must be avoided when applying LR to the analysis of behavioural phase state. Finally, I reject the idea that ‘standardised models’ provide a valid shortcut to estimating phase state across different developmental stages, strains or species. The points addressed here are pertinent to any research on transitions between complex phenotypes and behavioural syndromes.

## Introduction

A central question in animal behaviour is the extent to which individual differences in multiple behavioural traits are integrated together into behavioural syndromes (Sih et al., 2004; Dingemanse et al., 2010; Wolf and Weissing, 2012) or with other phenotypic dimensions such as morphology and physiology to form complex integrated phenotypes (Pigliucci, 2003; Murren, 2012; Armbruster et al., 2014; Kern et al., 2016). Phase change in locusts is a paradigmatic example of phenotypic plasticity and integration. Locusts can transform between two very distinct integrated phenotypes or phases in response to changes in population density (Pener and Simpson, 2009). This capacity for ‘phase change’ underpins the formation and breakup of swarms (Simpson and Sword, 2008; Cullen et al., 2017). Locust populations that have experienced low density conditions for several generations comprise individuals in the solitarious phase, which is characterised by cryptic colouration and a cryptic behavioural strategy that includes sparse locomotor activity with a crepuscular diel pattern and an aversion to conspecifics. Conversely, populations that have experienced high density conditions
for several generations comprise individuals in the gregarious phase, which shows distinct morphometric ratios, aposematic colouration, and a very different behavioural strategy that includes high levels of activity with a diurnal diel pattern and a propensity to be attracted towards conspecifics. The two phases also differ profoundly in metabolic and endocrine physiology and reproductive biology (Pener and Simpson, 2009).

Full phase transformation thus entails changes in many aspects of the phenotype that unfold over very different time scales: behavioural changes can occur within a few hours (Roessingh et al., 1993), whereas morphological changes can only occur over weeks or months, primarily as animals moult, with further changes accruing trans-generationally. Long-term phase state can be easily assessed by measuring morphological variables, and is therefore widely used in field surveys to inform locust control operations (Dirsh, 1951, 1953). The behavioural phenotype, however, cannot be inferred from morphometric measurements because it changes more quickly to reflect the recent history of population density. It also follows that mechanistic laboratory studies that target different aspects of phase change (behavioural, physiological, morphological) must each operate on an appropriate time scale. Most mechanistic studies to date have focussed on *behavioural phase change*, and have therefore operated on a time scale of hours or days (Anstey et al., 2009; Ma et al., 2011; Ott et al., 2012; Guo et al., 2015). In contrast, manipulations targeted at morphological phase traits would necessarily take weeks to manifest.

A trivially obvious prerequisite for any analysis of a specific phase trait is a meaningful measure of that trait. To this end, Martín-Blázquez and Bakkali (2017) have recently proposed standardised logistic regression (LR) models for estimating ‘the gregariousness level’ of individuals of the desert locust (*Schistocerca gregaria* Forskål) for adoption by the research community, together with suggestions for extending their approach to other locust species including the migratory locust (*Locusta migratoria* L.). I agree that standardised, accessible, open and transparent methods are needed (see also Cullen et al., 2017), but the models promoted by Martín-Blázquez and Bakkali (2017) are, regrettably, fundamentally flawed. The present paper is not intended as a comprehensive and detailed critique of the paper by Martín-Blázquez and Bakkali (2017); instead, I discuss specific conceptual and methodological shortcomings of that work that are germane to any research on transitions between complex phenotypes.

## Methods

The raw data included in the Supporting Information of Martín-Blázquez and Bakkali (2017) were analysed in R (RRID:SCR_001905) version 3.3.3 (R Core Team, 2016) and RStudio (RRID:SCR_000432) version 1.0.143 (RStudio Inc., Boston, MA) running under OS X El Capitan version 10.11.6 (Apple Inc., Cupertino, CA). LR models were fitted using the function *glm* from the built-in R package *core* to exactly replicate the analysis in Martín-Blázquez and Bakkali (2017). Additionally, LR models were fitted using the *lrm* function in package *rms*, version 5.1-1 (Harrell Jr, 2016), to obtain bootstrap-corrected values for two measures of model performance: (1) Somers’ *D*, a measure of the rank discrimination of the model; and (2) the intercept and slope of the calibration line, a measure of model calibration (Harrell Jr, 2001). Coefficients of variation were compared using the test of Feltz and Miller (1996) as implemented in the function *asymptotic_test* from the package *cvequality*, version 0.1.1 (Marwick and Krishnamoorthy, 2017). All analyses are documented in the Supplemental Materials of the present paper, which include a report in PDF format and the .Rmd source code that generates the report reproducibly from the raw data of Martín-Blázquez and Bakkali (2017).

### Phase transitions do not occur along a single latent axis

Martín-Blázquez and Bakkali (2017) introduce the notion of an overall ‘gregariousness level’ that encompasses all gregarious phase characteristics, be they molecular, behavioural, physiological, morphological or otherwise. I shall argue that this premise is conceptually misguided and of very limited value for any mechanistic analysis of phase change. In a series of influential papers, Simpson and colleagues introduced multivariate LR models to the analysis of behavioural phase change in the desert locust (Roessingh et al., 1993; Roessingh and Simpson, 1994; Simpson et al., 1999), and the approach has subsequently been extended to other locust species including *Locusta migratoria* (Guo et al., 2011; Ma et al., 2011) and *Chortoicetes terminifera* (Gray et al., 2009; Cullen et al., 2010, 2012). In all instances, the aim was to quantify *behavioural* gregariousness from multiple behavioural traits. Importantly, this approach assumes that phase-related behavioural traits change in concert during phase transformation, such that they can be interpreted as manifestations of a single latent *behavioural phase state*. In the desert locust, this assumption is not entirely uncontroversial (Tanaka and Nishide, 2013) but reasonably well-supported by experimental data (Rogers et al., 2014); in other species, the behavioural coherence during phase change would need to be explicitly tested.

The notion in Martín-Blázquez and Bakkali (2017) of a ‘gregariousness level’ that encompasses *all* gregarious phase traits is altogether different. They state that

> “In order to successfully test functionality of a gene or molecule, quantitative measurements of the level of gregariousness are needed. Currently no valid molecular marker is available, thus the assessment of the degree of locust gregariousness is based on mathematical modeling.” (Martín-Blázquez and Bakkali, 2017, abstract).

According to this notion, one is resorting to LR models only in lieu of a single reliable marker— which could, in principle, be any trait that differs between phases:

> “There is a plethora of […] potential indicators of the state of a locust population. Apart from their developmental, survival, reproductive, immunological, and physiological differences, [the two] phases also differ in their morphology […] and behavior.” (Martín-Blázquez and Bakkali, 2017, p. 21).

This presumes that all phase-related traits are tightly coupled at all times, so that any one can serve as a measure of the same latent ‘gregariousness.’ It is a ground truth, however, that phase transformation entails a *decoupling* of different phase-related traits: some behaviours change within hours, whereas morphological changes take several generations to fully manifest (Pener and Simpson, 2009).

Martín-Blázquez and Bakkali (2017) set out with a LR model that is based on a combination of morphometric and behavioural variables (model *‘Sg_extended’*), and a version of this model (*‘Sg_extended_corrected’*) is one of two put forward for adoption by the research community. To be clear, it is perfectly feasible to define a statistically sound LR model that incorporates morphometric, behavioural and other kinds of traits to predict the probability that a locust has a *longterm history* of high population density. For locusts that are undergoing phase transition, however, the ‘probability of gregariousness’ (*P_greg_*) predicted by a morphological-behavioural ‘hybrid’ model will be wholly uninformative. This becomes clear when one considers a long-term solitarious locust that has been crowded for one day. Its morphology will not have changed, but in important aspects of its behaviour, such as locomotor activity, it will already be comparable with long-term gregarious locusts (Roessingh and Simpson, 1994). A first problem is that including both morphometric and behavioural predictors into a multivariate LR model does not guarantee that they will have equal weight—in fact, this outcome is unlikely. The estimation of the weights (coefficients) of the predictors is entirely data-dependent: each estimate reflects the association of the predictor with phase after adjusting for the associations of all other predictors with phase and, importantly, the degree of collinearity among predictors, with all estimates subject to stochasticity in the data sample to which the model is fitted. One or more predictor variables will dominate the prediction, and the dominant predictor(s) may be behavioural or morphological depending on the specific combination of predictors included in the model and the sample that the model is fitted to. For a one-day crowded locust, models that combine morphological and behavioural variables may therefore predict any *P_greg_* value between near-zero (high probability of ‘solitariousness’) and near-one (high probability of ‘gregariousness’).

However, that entering predictors does not determine their weighting is almost a side point because, whatever the model prediction for our one-day crowded locust, it will in any case be inappropriate with respect to behaviour, morphology, or both. If the model predicts a *P_greg_* close to zero, it is obviously useless for detecting behavioural gregarisation. A value close to one would be entirely wrong as an estimate of morphometric phase state. Martín-Blázquez and Bakkali (2017) may have hoped for a value of around 0.5—the prediction from a model in which behavioural and morphometric variables have equal weight—but this value too would be discordant with phase state: our one-day crowded locust is intermediate *neither* in morphology (it still has completely solitarious morphology) *nor* in behaviour (it behaves gregariously). Overall phase state cannot be measured on a single latent axis but requires a multidimensional metric space—at least a plane spanned by behaviour and morphology if we consider only those two aspects and were to assume that they can each be sensibly collapsed onto a single axis. Even this assumption is too simplistic, however, because different aspects of morphological phase change are known to be mechanistically decoupled from each other. For example, different components of the gregarious colouration can be induced by separate sensory stimuli (Lester et al., 2005).

### Behavioural predictors should not be adjusted for body size

Martín-Blázquez and Bakkali (2017) furthermore argue strongly that ‘speed-related behavioural variables’ should be ‘normalised’ for body size, criticise previous work for not having done so, and advocate dividing the ‘speed-related variables’ by the length of the hind leg femur. Their argument is based on their reported correlations between ‘speed-related’ and morphometric variables (Table S5 in the Supporting Information of Martín-Blázquez and Bakkali, 2017). Here, it is necessary to distinguish between correlations *within* a phase and correlations *across* the two phases—a distinction that is not explicit in Martín-Blázquez and Bakkali (2017). Gregarious desert locusts have shorter hind femora than solitarious locusts (Dirsh, 1951, 1953), yet they walk faster, more frequently and spend more time walking, and consequently they cover more ground during the assay duration (Ellis and Pearce, 1962; Roessingh et al., 1993; Rogers et al., 2014). Clearly, leg length does not explain these phase differences in locomotion, and the apparent correlations with hind femur length across the two phases are expected to be negative. The locusts used by Martín-Blázquez and Bakkali (2017) are unusual in that they apparently have about 1.6 × *longer* hind legs in the gregarious phase (final instar nymphs; my analysis, Fig. S1 and Tables S5, S6 in my Supplemental Material). This may reflect a strain peculiarity that has not been reported in any other lab or wild strain, inappropriate husbandry, or a data labelling error; but whatever the cause it results in an unexpected *positive* across-phase correlation between *hind femur length* and *average speed* (my analysis; Spearman’s *ρ* = 0.336; *N* = 66, *S* = 31832, *P* = 0.00589; Fig. S2 in my Supplemental Material).

For correlations *within* a phase, I have in most instances been unable to replicate the results in Martín-Blázquez and Bakkali (2017) from the raw data provided in their Supporting Information (see Table S8 in my Supplemental Material). For example, their Table S5 gives the correlation between *average speed* and *hind femur length* in gregarious nymphs as *r* = 0.442 (*N* = 51, *P* = 0.00128); I obtained *r* = −0.0612 (*N* = 51, *t*_49_ = −0.429, *P* = 0.670), which indicates that leg length has a negligible effect on locomotor speed. Also, several correlation coefficients in Martín-Blázquez and Bakkali’s (2017) Table 2 do not match those in their more detailed Table S5 (*e.g.*, correlation between *hind femur length* and raw *erratic movement*, their Table 2: 0.215; their Table S5: −0.065; neither matches my calculation).

Nevertheless, one must concede that such correlations are plausible in principle. *If*, however, one were to commit to including both morphometric and behavioural predictors in a LR model, then adjusting upfront for a correlation between them is misguided because multivariate modelling already accounts for any correlations among predictors. While the individual associations of correlated predictors with the dependent variable cannot be resolved, such correlations do not affect predictive model performance (Harrell Jr, 2001). Martín-Blázquez and Bakkali’s (2017) misconception about how predictors are weighed in multivariate model fits is apparent from their statement,

> “To detect highly correlated variables that might reinforce or bias the model toward a particular trait, we carried out pairwise correlations [...]” (Martín-Blázquez and Bakkali, 2017, p.14):

Correlated predictors may lead to problems with model *estimation*, but they do not ‘reinforce’ the model *predictions* because their joint information enters the prediction only once.

If, on the other hand, one rejects the inclusion of morphometric variables in the model on the grounds developed in the previous section, then dividing behavioural variables by femur length or any other measure of body size is misguided because it re-introduces morphometric characters into the model ‘through the back door.’ Long-term solitarious locusts typically have longer legs. After 4–24 h of crowding, their locomotor characteristics will be virtually the same as those of long-term gregarious locusts but the legs will obviously be no shorter than before. If locomotor variables are divided by hind femur length, freshly gregarised locusts (with long legs) will yield values lower than those of gregarious locusts (with shorter legs) and will therefore be assigned an erroneously lower *P_greg_* in what is intended as an assessment of pure behavioural gregariousness.

Adjusting behaviour by morphology is ill-advised even if one considers only long-term solitarious and gregarious locusts. In Martín-Blázquez and Bakkali’s (2017) unusual data, where gregarious final instar nymphs have *longer* legs than their solitarious counterparts, a clear phase difference in *average speed* is obliterated after dividing by *hind femur length* (Fig. S3 and Table S9 in my Supplemental Material). This outcome is of course purely coincidental in the sense that the phase difference in *average speed* is not caused by the phase difference in leg length. But the consequence is that, in Martín-Blázquez and Bakkali’s (2017) data for final instar nymphs, raw *average speed* is a reasonably useful predictor of phase in a univariate LR model, whereas the corresponding model based on ‘normalised’ *average speed* is hardly better than random guessing (Table S10 in my Supplemental Material). Rather than improving predictive accuracy, as intended by Martín-Blázquez and Bakkali (2017), the ‘normalisation’ annihilates the predictive power of *average speed* in their data.

To prove that dividing speed-related variables by hind femur length successfully ‘homogenises’ the variance, Martín-Blázquez and Bakkali (2017) compared raw and ‘normalised’ variances using Bartlett’s tests, which are significant in all cases (Martín-Blázquez and Bakkali, 2017, Table 3). These tests are meaningless, however, because variance depends on the measurement scale, and division by femur length changes the dimension and scale for only the ‘normalised’ dataset. Bartlett’s test is for comparing variances between different groups of data measured on the same scale, and will trivially give a significant result when applied to two versions of the same set of data measured on two different scales—dividing any set of values *X* by any factor *s* > 1 will reduce the variance. The authors draw further erroneous conclusions from Bartlett’s *K*^2^:

> “It should be noted that if the animals are of similar size (the solitary samples used for this analysis), the normalization has a significant but clearly weaker effect (Table 3).” (Martín-Blázquez and Bakkali, 2017, p.14)

Scale-dependency aside, however, *K*^2^ also depends on the sample size: the much lower values of *K*^2^ reported in their solitarious locusts reflect the almost 6-fold smaller sample (adults and final instar nymphs combined: *N* = 28 solitarious *vs N* = 161 gregarious). An appropriate test would be comparing the *coefficient of variation c*_v_ = *σ/μ*, a scale- and dimension-independent measure of relative dispersion (Feltz and Miller, 1996). In the example of the average speeds of solitarious and gregarious nymphs, division by femur length does not appreciably reduce the dispersion in Martín-Blázquez and Bakkali’s (2017) data (raw *c*_v_ = 0.8227, ‘normalised’ *c*_v_ = 0.7774; *N* = 66 each for raw and ‘normalised’, Feltz-Miller statistic = 0.0915, *P* = 0.762). While some locomotion-related variables may conceivably correlate with body size, the evidence presented in Martín-Blázquez and Bakkali (2017) is fallacious and the proposed remedy creates a problem rather than solving one.

### Sample-size requirements for multivariate LR models

The most critical methodological failing of Martín-Blázquez and Bakkali (2017), however, concerns the specific LR models that they put forward, which are based on a sample of 51 gregarious and 15 solitarious *S. gregaria* nymphs. How many predictors a LR model can reasonably accommodate is limited by the number of observations in the smaller group (here, 15 solitarious locusts; Harrell Jr, 2001). The ratio of this ‘limiting sample size’ to the number of regression coefficients (excluding the intercept) is known as ‘events per variable’ (EPV; van Smeden et al., 2016). If EPV is low, the model will be unreliable; that is, it will not predict future observations as well as it appeared to predict on the present sample. Furthermore, there is an increased likelihood of ‘complete separation’ of the two groups (here, of the two phases), in which case the model estimation fails altogether. The two models advocated in Martín-Blázquez and Bakkali (2017), *‘Sg_extended / Sg_extended_corrected’* and *‘Sg_non-morphometric’*, have 13 and 10 predictors, respectively, which with 15 solitarious locusts means less than 2 EPV. Between 10 and 20 EPV are widely considered a minimum (Harrell Jr, 2001), and while this is no hard and fast rule (van Smeden et al., 2016), attempting to fit LR models with less than 2 EPV is hopeless.

This is demonstrated very instructively by inspection of the model fitting results reported by Martín-Blázquez and Bakkali (2017). Replicating the LR fits in the same software as used in the paper (R, *glm* function) reproduces the numerical results with two-digit accuracy or better (Tables S13, S15, S20 in my Supplemental Material). The slight discrepancies likely reflect platform differences in floating point arithmetic that only manifest when models are poorly estimable. Large apparent discrepancies are resolved as manuscript errors in Martín-Blázquez and Bakkali (2017): First, in their Table 5, some variable names are switched: *average acceleration* should read *stop ratio; stop ratio* should read *turn ratio;* and *turn ratio* should read *average acceleration* (cf. their Table S5, where the labels are correct). Second, for models *‘Sg_extended’* and *‘Sg_non-morphometric’*, all coefficients and standard errors (SE) reported in the paper (Martín-Blázquez and Bakkali, 2017, Tables 5 and S5) are 10^3^ too high (Tables S13, S15 in my Supplemental Material). This systematic error nevertheless accounts only in part for the unreasonably large coefficients and SEs reported for the two advocated models. Another contributing factor is the extreme scaling of the ‘normalised’ speed-related variables (means between about 2 × 10^−4^and 2 − 10^4^; Table S4 in my Supplemental Material), which makes it hard to spot conspicuously large SEs that are indicative of collinearity problems.

After means-centring and scaling to *σ* = 1, refitting *‘Sg_non-morphometric’* flags up pathologically large SE estimates for *average speed* (*β_as_* = −3.25, SE = 75.9, *z* = −0.043, *P* = 0.966) and *average acceleration* (*β_aa_* = 2.87, SE = 75.8, *z* = 0.038, *P* = 0.970; Table S16 in my Supplemental Material) that can be traced to near-perfect collinearity between them (*r* = 0.9998; Fig. S4 in my Supplemental Material). This indicates a fundamental error in the measurement or calculation of one or both variables. More generally, however, the example highlights the importance of examining whether the estimated coefficients that one has obtained in the model fitting are *sensible.* At face value, *β_as_* = −3.25 would mean locusts that have a higher *average speed* are more likely to be *solitarious*, which would be very unexpected; but since in Martín-Blázquez and Bakkali’s (2017) data *average acceleration* is near-perfectly collinear with *average speed* and *β_aa_* = 2.87, the two effectively cancel out. Furthermore, *final choice* (the side of the arena where the locust was at the end of the assay) encodes the sign of *last coordinate* in the arena, which makes *final choice* redundant. This resulted in a model fit where the coefficient is positive for *final choice*, but *negative* for *last coordinate* (although not significantly different from zero; Table S16 in my Supplemental Material). It is thus important to examine whether the directions (signs) of the estimated coefficients are consistent both internally and with prior subject knowledge; where they are not, the model may be ill-specified.

After removing *average acceleration* and *final choice*, the fit and predictive performance of the *‘Sg_non-morphometric’* model can be validated by bootstrapping, although model fitting still fails in about 15% of bootstrap samples due to divergence or singularity (*B* = 1000 bootstrap samples; Table S19 in my Supplemental Material). The results indicate that the model’s rank-discrimination is mediocre (bias-corrected Somers’s *D* = 0.66, where 0 is no predictive power, 1 is perfect), but more importantly that the calibration is very poor: the bias-corrected calibration line has an intercept of 0.49 (where 0 is perfect, 1 is worst) and a slope of 0.45 (where 1 is perfect, 0 is worst), demonstrating the extreme overfitting that occurs with less than 2 EPV.

For the *‘Sg_extended’* / *‘Sg_extended_corrected’* model, *all* the coefficients are extremely large even after rescaling the predictors to *σ* =1, the SEs are astronomical and, consequently, all associated *P* values exceed 0.99 (Tables S13, S14 in my Supplemental Material), as they do in (Martín-Blázquez and Bakkali, 2017, Table S5 in their Supporting Information). Such evidently nonsensical results are symptoms of a failed model fit. Replicating exactly the analysis in the paper shows that the *glm* function issues two warning messages when fitting *‘Sg_extended’*: *‘glm.fit: algorithm did not converge’* and *‘fitted probabilities numerically 0 or 1 occurred’* which together indicate that complete separation has occurred (see p. 11 in my Supplemental Material). Martín-Blázquez and Bakkali (2017) misinterpreted complete separation as excellent predictive accuracy—they describe their model as highly accurate because it *‘detected all the 51 gregarious nymphs as gregarious with 100% probabilities and attributed 0% gregariousness probability to all our 15 solitary nymphs’.* The authors also considered a five-predictor model (*‘Sg_low-redundancy’*) which, while still hopelessly underpowered (3 EPV), did not result in complete separation. Of the three models in Martín-Blázquez and Bakkali (2017), *‘Sg_low-redundancy’* is the least deficient from a purely technical point of view. It has reasonable rank-discrimination (bootstrap bias-corrected Somers’ *D* = 0.85) and shows considerable but tolerable overfitting (bootstrap bias-corrected linear calibration intercept = 0.09, slope = 0.80)—although the model fit fails in about 20% of bootstrap samples (Tables S20–S22 in my Supplemental Material). Thinking wrongly that this model *‘is not as accurate in predicting gregarious locusts as the ‘Sg_extended’ model,’* the authors discarded it in favour of the fatally flawed *‘Sg_extended’* model. In attempting to validate *‘Sg_extended’* on data from locusts reared at a range of intermediate population densities, Martín-Blázquez and Bakkali (2017) found that it predicted extremely dichotomised probabilities that did not match the expected intermediate phase states. This inevitable consequence of the failed model fit cannot be ‘fixed’ by shrinking the logit predictions ad hoc by an arbitrary ‘homogeneous correction’ factor, which is what Martín-Blázquez and Bakkali (2017) did to arrive at their final *‘Sg_extended_corrected’* model that they recommended for uptake by the research community. Shrinkage methods that are grounded in statistical theory such as ridge and LASSO regression are available for handling the ‘many predictors / few samples’ problem (Hastie et al., 2009). Even these methods, however, cannot overcome the limitation inherent to a sample size of 15, because nothing can generate information beyond what is provided by the sample.

### Different stadia, strains and species need different models

Previous studies have used LR models that were based on samples from the same species, strain and developmental stage as those locust that the model was used to predict on. Studies on first-instar, final-instar or adult *S. gregaria* used models fitted to first- or final-instar nymphs or adults, respectively, of the same laboratory strain of that species (Roessingh et al., 1993; Islam et al., 1994; Bouaïchi et al., 1995); studies on different instars of *L. migratoria* used *L. migratoria* instar-specific models (Ma et al., 2011). The ultimate aim of Martín-Blázquez and Bakkali (2017) are ‘standardised’ models that can be applied across different laboratory strains, and ideally across different stadia and species (in their study, final instar nymphs and adults of *S. gregaria* and *L. migratoria*). Martín-Blázquez and Bakkali (2017) gave explanations for why their efforts were only partially successful by their own lights. The validity of these explanations is limited by their small sample sizes and by fundamental statistical misconceptions that I have discussed above. Here I consider the broader question whether ‘standardised models’ are a sound proposition in principle.

There are two distinct, although connected, aspects of ‘standardisation’ to Martín-Blázquez and Bakkali’s (2017) models. First, they are intended as community standards; second, they are based on behavioural predictors that have been ‘standardised’ for morphometric differences. This second aspect was at least in part motivated by the hope that it would increase the validity of the models across different developmental stages, strains and species. Although Martín-Blázquez and Bakkali (2017) concluded that their *S. gregaria* models did not perform well in *L. migratoria*, they recommended their *‘Sg_non-morphometric’* model, which was fitted to final instars, for use in adults:

> “For testing adults or the same nymphs at different time points (if they do not molt), we suggest using the ‘Sg_non-morphometric’ model (that does not include morphometric variables).” (p. 23).

I refer to my earlier argument for why this model, although not containing overt morphometric variables, is nevertheless contaminated with morphometric information. Above, I have shown that ‘standardising’ behaviour by morphology is inappropriate if the locusts that the model is fitted to are of the same developmental stage as those that the model is then used to predict on. ‘Standardising’ behaviour by morphology is even more inappropriate if the model is then used to predict on locusts of a different developmental stage. This is readily seen when plotting Martín-Blázquez and Bakkali’s (2017) data for *average speed* over *hind femur length* for gregarious nymphs and adults: adults have distinctly longer hind femora (the distributions barely overlap), but their *average speeds* are *lower* in Martín-Blázquez and Bakkali’s (2017) data (Fig. S5 and Table S23 in my Supplemental Material). Consequently, dividing *average speed* by *femur length* increases, rather than reduces, the (now no longer purely behavioural) difference in ‘standardised average speed’ between gregarious nymphs and adults (Fig. S6 in my Supplemental Material). Because behavioural differences between developmental stages, strains and species are not merely caused by morphology, they cannot be made disappear by dividing behavioural variables by femur length, or by any other morphometric.

Are standardised models without inappropriate ‘morphometric standardisation’ of behavioural predictors a valid proposition? Martín-Blázquez and Bakkali (2017) give two motivations for their drive towards standardised models. First, it would do away with the need for each research group having to build their own model. This would save a great deal of effort, because building a model requires observations from a large number of ‘reference locusts’ (certainly more than the 15 solitarious locusts used by Martín-Blázquez and Bakkali, 2017). Second, the use of a common model would facilitate direct comparisons between results from different research groups.

At first, these arguments may appear attractive or even compelling. Their fallacy becomes apparent upon reflection on what we ask a standard model to do. We ask it to spare us the effort of collecting data for a different developmental stage, strain or species of locust; but these are the very data that are needed to ascertain the validity of the ‘standardised model’ in that developmental stage, strain or species. One does not know a priori that a ‘standardised model’ based on a sample of locusts from, e.g., strain A will adequately predict on locusts from a different strain B. Whether it does cannot be ascertained by validating the model in just a handful of locusts from B. To assess whether strain differences can be safely ignored, one needs an adequate number of observations on both phases from both strains under comparable laboratory conditions; fit a model that includes strain as an additional predictor variable together with all interactions between strain and the remaining predictors (Model 1); and fit the alternative model that does not include strain or any of its interactions (Model 2). In the symbolic form used to specify models in R, the models for *k* predictors *x*_1_, … *x_k_* would be:

Model 1 : *phase* ∼ (*x*_1_ + *x*_2_ … + *x_k_*) * *strain*

Model 2 : *phase* ∼ *x*_1_ + *x*_2_ … + *x_k_*

One can then compare the two models based on, e.g., AIC or a likelihood ratio test. Only if in an *adequately powered* analysis AIC were lower for Model 2 or if the test came out decisively ‘non-significant’ (e.g., *P* > 0.15) would we accept Model 2 as likely to predict well in both strains. Key here is *adequately powered:* to decide whether Model 2 is adequate, we need to have adequate data on both strains in the first place. This reveals the fallacy that a ‘standardised model’ can save us the effort of collecting an adequately large sample in the strain that we want to predict on. The full model needs 2*k* + 2 coefficients to be estimated (including the intercept), and by rule of thumb the minimum sample size would be about 15(2*k* + 2) locusts of each phase, balanced with respect to strain. A strain-specific model requires *k* + 1 coefficients to be estimated, and the minimum sample size would be about 15(*k* + 1) locusts of each phase, of that strain. The conclusion from this is that validating an existing model in a new strain needs about as many locusts as building a new model.

Thus, a final and fundamental objection to the ‘standardised models’ suggested by Martín-Blázquez and Bakkali (2017) concerns the very proposition that one and the same model fit can be meaningfully generalised to different laboratory strains, let alone different developmental stages or species. This proposition fails to distinguish between a standardised *assay* and a standardised *fitted model.* There is a case for standardising the assay conditions, although what aspects to standardise needs discussion—a behavioural arena for adults needs to be larger than one for first instar nymphs of the same species. The research community should agree on a standard set of predictor variables for LR models of behavioural phase state, and on transparent algorithms that define these variables. But the *fitted model* cannot be simply transplanted from one locust species, strain or developmental stage to another.

## Conclusions

Although I appreciate Martín-Blázquez and Bakkali’s (2017) effort to promote open and transparent research, the advocated statistical proce-dures and models are conceptually and methodologically flawed and should not be adopted. The premise of an overall ‘level of gregariousness’ that conflates morphological and behavioural phase characteristics is conceptually misguided, and statistical models based on this premise are of extremely limited use for the analysis of phase transitions. Standardising behaviour by body size creates a problem rather than solving one. Well-calibrated multivariate regression models require adequately large samples of locusts drawn from the population that one is working with, and the proposition that ‘standardised’ models offer a shortcut to models that are valid across different developmental stages, strains or species needs to be rejected.

## Acknowledgments

SRO thanks Dr Tom Matheson (University of Leicester) and Dr Stephen M. Rogers (University of Cambridge) for helpful discussions.

## Author Contributions

SRO is the sole author of this article. SRO conceived this article and the arguments that are put forward in it; carried out the statistical analyses, as documented in the supplemental materials; and wrote the manuscript.

## Supplemental Material

All analyses are documented in the Supplemental Material of the present paper, which includes a report in PDF format and the .Rmd source code that generates the report reproducibly from the raw data of Martín-Blázquez and Bakkali (2017).

## Data Availability

The datasets analysed for this study can be found in the Supporting Information of Martín-Blázquez and Bakkali (2017, DOI: 10.1111/eea.12564). The exact version of these data that was used as input for the analyses in the present study is included as Supplemental Data with this present paper.

